# Long Range PCR-based deep sequencing for haplotype determination in mixed HCMV infections

**DOI:** 10.1101/2021.07.05.451103

**Authors:** Nadja Brait, Büşra Külekçi, Irene Goerzer

**Author notes:** The authors are joint authors and contributed equally to this study (Nadja Brait, Büşra Külekçi). Groningen Institute for Evolutionary Life Sciences, University of Groningen, Groningen, Netherlands (Nadja Brait). **Repositories:** Sequence data generated in this study were deposited in GenBank with the accession numbers MW560357-MW560373. Raw data of Illumina and PacBio sequencing were submitted to the NCBI Sequence Read Archive (SRA) under BioProject ID PRJNA698084 and BioSample accession numbers SAMN17705199-SAMN17705210, SAMN17705219-SAMN17705248.

## Abstract

Short read sequencing, which has extensively been used to decipher the genome diversity of human cytomegalovirus (HCMV) strains, often falls short to assess co-linearity of non-adjacent polymorphic sites in mixed HCMV populations. In the present study, we established a long amplicon sequencing workflow to identify number and relative quantities of unique HCMV haplotypes in mixtures. Accordingly, long read PacBio sequencing was applied to amplicons spanning over multiple polymorphic sites. Initial validation of this approach was performed with defined HCMV DNA templates derived from cell-free viruses and was further tested for its suitability on patient samples carrying mixed HCMV infections.

Our data show that artificial HCMV DNA mixtures were correctly determined upon long amplicon sequencing down to 1% abundance of the minor DNA source. Total error rate of mapped reads ranged from 0.17 to 0.43 depending on the stringency of quality trimming. PCR products of up to 7.7 kb and a GC content <55% were efficiently generated when DNA was directly isolated from bronchoalveolar lavage samples, yet long range PCR may display a slightly lower sensitivity compared to short amplicons. In a single sample, up to three distinct haplotypes were identified showing varying relative frequencies. Intra-patient haplotype diversity is unevenly distributed across the target site and often interspersed by long identical stretches, thus unable to be linked by short reads. Moreover, diversity at single polymorphic regions as assessed by short amplicon sequencing may markedly underestimate the overall diversity of mixed populations.

Quantitative haplotype determination by long amplicon sequencing provides a novel approach for HCMV strain characterisation in mixed infected samples which can be scaled up to cover the majority of the genome. This will substantially improve our understanding of intra-host HCMV strain diversity and its dynamic behaviour.

**Impact statement:** Human cytomegalovirus (HCMV), a large enveloped DNA virus, displays the highest inter-host genome variability among all human herpesviruses. Primary infection, reinfection and reactivation are mostly asymptomatic but may cause devastating harm in congenitally infected newborns and in immunosuppressed individuals. Multiple distinct strains circulate in humans, each characterised by a unique assembly of well-defined polymorphic genes, most of which are linked to cell entry, persistence and immune evasion. Mixed HCMV strain infections are common and may pose a high pathogenic potential for patients at risk for symptomatic infections. To better understand the biological behaviour and dynamics of individual viral genomes it is inevitable to assess the co-linearity of polymorphic sites in a genetically heterogeneous population. In this study, we established and successfully applied a long read sequencing technique to long amplicons and identified co-linear genome stretches (haplotypes) in patient samples with mixed HCMV populations. This strategy for haplotype determination allows linkage analysis of multiple non-adjacent polymorphic sites along up to 7.7 kb. This allows a better approximation to the true strain diversity in mixed samples, which short read sequencing approaches failed to do. Thereby, improving our knowledge on mixed HCMV infections important for the clinical outcome, diagnostics, treatment and vaccine development.

**Data Summary:** Sequence data generated in this study were deposited in GenBank with the accession numbers MW560357-MW560373. Raw data of Illumina and PacBio sequencing were submitted to the NCBI Sequence Read Archive (SRA) under project number SUB8972240. BioSample accession numbers are provided in Supplementary Table 3 and 4.

Additional sequence data for reference purposes were accessed from GenBank. Accession numbers are listed in Supplementary Table 6 and 7.

## Introduction

It is well-known that human cytomegalovirus (HCMV) seropositive persons can be infected with more than one HCMV strain, either synchronously or sequentially [1-6]. Solid organ transplant patients, in particular, are at high risk for harbouring complex HCMV strain mixtures when multiple donor HCMV strains are transmitted to an already HCMV seropositive recipient probably associated with a poorer clinical outcome [7-14].

The genome sequence of an individual HCMV strain is characterised by a unique combination of linked sequence patterns of all polymorphic regions along the whole genome [15]. Individual polymorphic sites within the HCMV genome, either comprising a complete gene or a highly variable section thereof, have been widely investigated with first and second generation short read sequencing [16, 17]. These investigations uniformly show that almost all polymorphic regions cluster into defined genotypes [18, 19]. This allows to assign even short sequence reads to reference genotype sequences by the use of genotype-defining signature sequence patterns of the respective polymorphic regions [10, 19, 20]. In recent years, widely used in-depth short read sequencing of the whole genome comprehensively assessed genome-wide intra- and inter-host diversity at the nucleotide level, and intra-host variability over time [18, 19, 21-25]. Moreover, these studies revealed that recombination between and within polymorphic regions of different HCMV strains have been common and single nucleotide variants that may emerge during replication periods will further contribute to the overall strain diversity. Accordingly, it appears that a huge variety of different HCMV strains circulate in the human population and the frequency of occurrence of completely identical “strain genotypes” among HCMV-infected persons seems to be very rare [19].

Due to short read lengths of second generation sequencing techniques, determination of individual genomes within mixtures can be challenging. It requires model-based analysis to reconstruct unique haplotypes from short sequencing reads [19, 22, 25, 26] but this might be difficult for genomic segments interspersed by longer conserved sections, low-complexity repetitions and GC-deviant regions [27].

In contrast to short read sequencing platforms, single molecule real-time (SMRT) sequencing from Pacific Biosciences (PacBio) and MinION Nanopore sequencing from Oxford Nanopore Technologies exceed with their ability to perform novel genome analyses, due to their extended read length [28, 29]. Unlike their predecessor, third generation single-molecule technologies can frequently generate read lengths up to 30 kb and have been used to fill many gaps in reference genomes, which could not be covered due to short read lengths [30, 31].

Still, there is limited data on the number of unique HCMV DNA molecules within mixed HCMV populations, which in the following will be termed HCMV haplotypes. It is still unknown how specific combinations of polymorphic regions present on an individual HCMV DNA molecule affect pathogenicity or replication efficiency.

In the present study we assessed how enrichment of HCMV DNA by long range PCR and long read sequencing enables the identification of linkages between non-adjacent polymorphic regions present on single HCMV DNA molecules. The newly developed protocol was successfully applied to a selection of bronchoalveolar lavage samples (BALs) displaying mixed HCMV infections. This approach provides novel insights into the composition of individual haplotypes within a mixed HCMV strain population.

## Methods

### Primer design for long range PCR

Primers are depicted in Supplementary Table 1. A total of 310 whole genome sequences were screened for conserved regions in the NCBI Multiple Sequence Alignment Viewer 1.11.1. Optimal primer sequences within the conserved regions were determined with Primer-Blast. In silico primer design was performed using the GeneArt® Primer and Construct Design Tool with the Single Site-Directed Mutagenesis option. Sequences of desired target regions which include highly polymorphic regions within the UL section were derived from strain Merlin (AY446894.2) and a blastn search with the Nucleotide collection database (nr/nt) was performed. Single CDS, partial and artificial genomes were excluded.

### Preparation of HCMV DNA used for sequencing

#### Bacterial artificial chromosomes (BAC) HCMV DNA from E.coli

TB40-BAC4-DNA was purified from 400 ml of *E. coli* overnight culture using the Nucleobond BAC 100 kit (Machery-Nagel) for low-copy plasmid purification. All steps were done according to the manufacturer. Purified BAC-DNA was eluted in 50 µl nuclease-free deionized water, quantified using Nanodrop and stored at 4°C and under no conditions frozen to avoid DNA fragmentation. Purified BAC-DNA was used to establish long range PCR assays.

#### HCMV DNA from fibroblast supernatant

Cryopreserved virus-stock vials of strains Merlin (AY446894) and TB40-BAC4-luc (derivative of TB-BAC4, EF999921 [32]) derived from human foreskin fibroblast (HFF) supernatant were used for HCMV-DNA purification. Therefore, 500 µl of thawed virus stock samples were transferred into 2 ml of lysis buffer and eluted in 50 μl elution buffer using the bead-based NucliSens EasyMagextractor (BioMérieux). Quantification of HCMV-DNA was done by HCMV-specific qPCR (see below) and stored at 4°C before being directly subjected to Illumina and PacBio sequencing or taken as template for long range PCR.

#### HCMV DNA from BAL samples

Six BAL samples stored at -20°C, all from patients who received lung transplants at the Medical University of Vienna between 2014 and 2016, were investigated. DNA was isolated using the QIAamp Viral RNA Mini kit (Qiagen). Therefore, 250 µl of BAL solution was lysed and further purified as described in the manufacturer’s protocol. DNA was eluted from columns with 70 µl elution buffer. HCMV DNA was quantified by HCMV-specific qPCR (see below) and stored at 4°C before being subjected to short and long range PCR.

#### Artificial mixtures of two distinct HCMV-DNA genomes

HCMV-DNA either purified from Merlin or TB40-BAC4-luc virus stocks were diluted to the appropriate concentrations and mixed to achieve the ratios as listed in Table 5. Five µl of each mixture (total HCMV-DNA ranged from 3×10^6^ to 9×10^6^ copies per reaction) was used as template DNA for amplification by long range PCR.

### DNA Quantification

Purity and content of BAC-DNA and cell-culture-derived DNA was quantified using the NanoDrop 1000 tool (Peqlab). Amount of PCR amplicons was determined using the Qubit™ double-stranded DNA High-Sensitivity Assay according to the manufacturer’s instructions (Thermo Fisher Scientific) on the Qubit 2.0 fluorometer. HCMV-specific DNA of cell-culture-derived DNA and BAL samples was quantified by an in-house qPCR as previously described [33] and cell-culture-derived HCMV DNA for generation of artificial mixtures was additionally quantified by a gH genotype-specific PCR, also as previously described [34].

### Long range PCR amplification

PCR enzymes initially used for evaluation of sensitivity and specificity of long range PCR are listed in Table 1 and Supplementary Table 2. Target regions chosen for evaluation are shown in Figure 1. After evaluation, LA Taq HS DNA polymerase kit from TaKaRa (TaKaRa Bio) conveyed the best performance and was therefore further used. For long range PCR 15µl mastermix (0.25 µl Taq, 2.5 µl 10x buffer, 4 µl dNTPs, 9.75 µl nuclease-free water, 1.25 primer each) was combined with 10 µl extracted DNA. Correlating copy number input for individual samples can be seen in Tables 4a and 4b and Supplementary Table 11. The cycling program was 94°C for 1 min, 30x (98°C for 10 sec, 68°C for 1 min/kb) and without an additional extension step to avoid PCR-mediated recombination. Small aliquots of the PCR products were visualised on analytical agarose gels, then quantified by Qubit and subjected to library preparation for Illumina and PacBio sequencing.

**Table 1.** Efficiency of long range PCR for the distinct fragments by the use of three commercially available enzymes.

| Amplicon | Fragment length<br>in bases | GC content in<br>% | total BAC-DNA as template (100ng) |  |  | cell-culture derived DNA as template* |  |
| --- | --- | --- | --- | --- | --- | --- | --- |
|  |  |  | TaK | Pro | Qia | TaK | TaK |
|  |  |  | efficiency | efficiency | efficiency | specificity (1x10<sup>5</sup><br>copies/reaction) | sensitivity<br>(copies/reaction) |
| F1.1 | 7807 | 56.5 | high | low | low | low | 300 |
| F1.2 | 13807 | 58.9 | high | - | - | - | nd |
| F1.3 | 18015 | 56.5 | - | - | - | nd | nd |
| F2.1 | 8278 | 57.1 | high | - | - | - | nd |
| F2.2 | 13348 | 58.8 | high | - | - | - | nd |
| F2.3 | 17861 | 56.4 | - | - | - | nd | nd |
| F3 | 7705 | 53.7 | high | high | low | high | 30 |
| F4 | 6671 | 48.5 | high | nd | nd | high | 30 |

\* two different HCMV strains (TB40E and Merlin); high and low efficiency as determined by the strength of visible bands of the correct length; high specificity as determined by the absence of unspecific bands; TaK: LA Taq Hot Start Version Polymerase; Qia: LongRange PCR Polymerase; Pro: Promega Go Taq Long; nd: not determined; "-": no visible band or smear.

**Figure 1:**
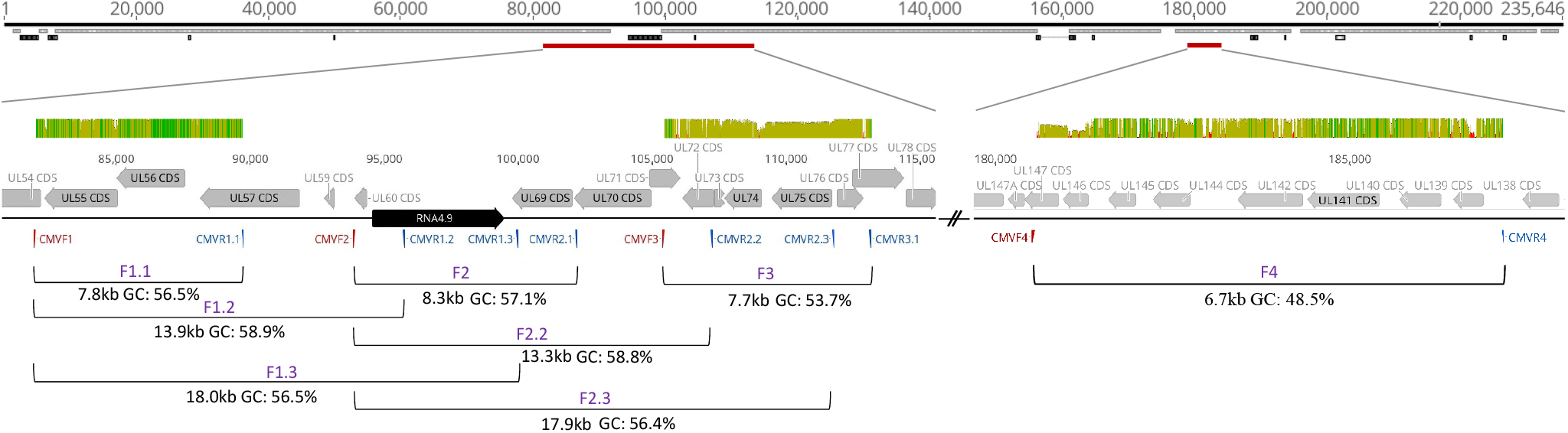
Schematic representation of target sequences along a HCMV genome. Shown is an overview of polymorphic amplified regions and their genetic diversity along the HCMV genome strain Merlin (AY446894). The figure consists of four separate tracks. In the top track, a mini-map of the genome with a total bases of 235,646 is shown. Annotated CDS and non-coding RNAs are depicted in grey and black, respectively. Target regions for amplicon generation are depicted in red and are described in detail in track 2-4. For track 2, defined polymorphic regions were put through a blastn search. Over 200 published haplotypes for the described regions were aligned. The mean pairwise identity over all pairs in the column is depicted. Green: 100% identity, greenish-brown: at least 30% identity but less than 100%, red: less than 30%. Displayed above the sequences (3^rd^ track), grey and black arrows again represent CDS and non-coding RNAs, respectively. Forward and reverse primer (red and blue) are depicted below the sequence. The bottom track indicates individual fragment sizes and GC content of eight polymorphic target regions. Annotations were generated with geneious prime 2019.0.3.

### Short range PCR amplification

HCMV-DNA positive BAL samples were screened for mixed genotype infections by short range PCR amplification of polymorphic target regions within envelope glycoprotein N (gN), envelope glycoprotein O (gO), and UL146 (nested PCR). Detailed description of PCR amplicons, associated primers and respective annealing temperatures are listed in Supplementary Table 1. For PCR, 19 µl AmpliTaq Gold 360 Mastermix (Applied Biosystems) and 0.5 µl of each primer was combined with 5 µl of extracted BAL DNA. Initial denaturation was performed at 95°C for 10 min followed by 40 cycles of 95°C for 1 min, annealing for 1 min at 50-61°C, and elongation at 72° for 1 min, with a final extension time of 5 min at 72°. For the second step of the nested UL146 PCR, 40 µl master mix and 8 µl PCR grade water was mixed with 2 µl of the first PCR product. Small aliquots of the PCR products were visualised on analytical agarose gels, then quantified by Qubit and subjected to library preparation for Illumina sequencing.

### PacBio SMRT sequencing

SMRT bell library preparation and sequencing were performed by the Next Generation Sequencing Facility at Vienna BioCenter Core Facilities. Long range PCR amplicons and non-enriched DNA were further purified using the QIAEX II Gel extraction Kit (Qiagen). Due to indispensable loss of HCMV-DNA (>80%) during gel extraction, both non-enriched DNA and PCR products were subjected to the protocol: Desalting and Concentrating DNA Solutions. All steps were performed as advertised in the protocol and the HCMV-DNA was eluted in 20 µl of Tris buffer. For library preparation a minimum DNA input of 700 ng and 2.5 µg for amplicon and whole genome sequencing was required, respectively. Initial quantification and purity were measured with the NanoDrop 1000 tool (Peqlab) and a Qubit 2.0 fluorometer (Thermo Fisher). Large fragment analysis was performed with a Femto Pulse system and the genomic DNA 165 kb Kit (Agilent). Non-enriched DNA was sheared to an expected fragment length of <10 kb. Samples were indexed and multiplexed according to the Sequencing facility. Subsequent to library preparation a blue pippin size selection (Sage Science Inc.) was used to isolate target fragment lengths of long range PCR amplicons. Sequencing was performed on a PacBio-Sequel system for 10 and 20 hours for cell-culture derived and BAL samples, respectively. Details on PacBio sequencing run information is listed in Supplementary Table 3.

### Illumina Sequencing

Two ng input DNA per sample was used for library preparation using the Nextera XT library preparation kit, and samples were indexed using the Nextera XT index kit (Illumina). After index PCR, samples were purified with 45 µl of Agencourt AMPure XP magnetic beads with a sample to beads ratio of 3:2 (Beckman Coulter), normalized by Qubit quantification and pooled to generate a 4 nM library. To compensate for low diversity libraries, a 12 pM PhiX control spike-in of 2.5% was added. Single and paired-end sequencing (150 to 250 cycles, V2 kits) was done on an Illumina MiSeq instrument with automatic adapter trimming (Illumina). Raw reads in fastq format that passed filters were used for analysis using CLC Genomics Workbench 12.0 (Qiagen). Detailed information on MiSeq sequencing runs is listed in Supplementary Table 4.

### Bioinformatical workflows for PacBio and Illumina reads

#### Generation of circular consensus sequence reads upon PacBio sequencing

Bam files of PacBio raw reads were demultiplexed using Lima (https://github.com/pacificbiosciences/barcoding/) with default parameters and *symmetric options*. In order to generate Highly Accurate Single-Molecule Consensus Reads, demultiplexed subreads were aligned with the PacBio circular consensus sequence (ccs) tool of Bioconda (https://github.com/PacificBiosciences/ccs). No full-length subreads were required for ccs generation and a minimum predicted accuracy of 0 was chosen. To prevent heteroduplexes, consensus sequences for each strand of a molecule were separated by strand. Resulting ccs reads were used for further analysis using CLC Genomics Workbench 12.0 (Qiagen Bioinformatics).

#### Trimming and mapping for error calculation

Illumina fastq reads were imported as paired-end reads and PacBio ccs were imported as single reads into CLC Genomics Workbench. Raw reads were quality trimmed by using the modified-Mott trimming algorithm 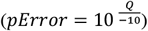 with a base-calling error probability of 0.01 or 0.001 (Phred quality score of 20 or 30, respectively), an ambiguous limit of 2 in a read, a minimum and maximum number of nucleotides in reads of 100 and 9000, respectively. Human genomic DNA reads were filtered out by randomly mapping them against the latest reference genome GRCh38 (accession GCA_000001405.28), with default setting parameters for match/mismatch scores and insertion/deletion costs, length fraction of 0.3, and similarity fraction of 0.8. PacBio-derived reads were further trimmed to exclude reads lower 500 nucleotides. Remaining reads from both sequencing platforms were randomly mapped to the pathogen reference sequences Merlin and TB40-BAC4-luc, respectively, with default setting parameters for match/mismatch scores and insertion/deletion costs, length fraction of 0.3, and similarity fraction of 0.8. Details are listed in Table 2 and Supplementary Table 5.

**Table 2.** Trimming, filtering, and mapping of PacBio-derived reads for error estimation with a base-calling error probability of 0.01 and 0.001.

| pError | Template HCMV DNA | Input DNA | Reference length in kb | Number and average read length of sequencing reads |  |  |  |  |  |  |  |  |
| --- | --- | --- | --- | --- | --- | --- | --- | --- | --- | --- | --- | --- |
|  |  |  |  |  | raw reads | after qtrim | after hu removal | % of reads <500 bases | after ltrim | % HCMV mapping | map to reference | average coverage |
| **0.01** | Merlin | amplicons | 15,5 | number, % | 92128 | 63604 | 30632 | 18,7 | 24909 | 98,26% | 24475 | 6885 |
|  |  |  |  | length | 4656 | 2820 | 3741 |  | 4547 |  |  |  |
|  | TB40-BAC4-luc | amplicons | 15,5 | number, % | 73439 | 46547 | 42060 | 19,1 | 34041 | 99,53% | 33881 | 9062 |
|  |  |  |  | length | 6162 | 3590 | 3767 |  | 4601 |  |  |  |
|  | Merlin | non-enriched | 237 | number, % | 143728 | 102444 | 5166 | 13,9 | 4447 | 85,29% | 3793 | 45 |
|  |  |  |  | length | 4378 | 2722 | 2956 |  | 3397 |  |  |  |
|  | TB40-BAC4-luc | non-enriched | 235 | number, % | 96966 | 72804 | 9122 | 12,6 | 7975 | 93,63% | 7467 | 73 |
|  |  |  |  | length | 3583 | 2439 | 2417 |  | 2731 |  |  |  |
| **0.001** | Merlin | amplicons | 15,5 | number, % | 92128 | 50031 | 22247 | 36,4 | 14154 | 97,76% | 13837 | 2444 |
|  |  |  |  | length | 4656 | 1631 | 1896 |  | 2843 |  |  |  |
|  | TB40-BAC4-luc | amplicons | 15,5 | number, % | 73439 | 34137 | 30361 | 37,2 | 19070 | 99,26% | 18929 | 3118 |
|  |  |  |  | length | 6162 | 1814 | 1865 |  | 2828 |  |  |  |
|  | Merlin | non-enriched | 237 | number, % | 143728 | 83991 | 4028 | 27,2 | 2931 | 86,49% | 2535 | 25 |
|  |  |  |  | length | 4378 | 1849 | 1934 |  | 2566 |  |  |  |
|  | TB40-BAC4-luc | non-enriched | 235 | number, % | 96966 | 60539 | 7402 | 23,2 | 5688 | 94,57% | 5379 | 45 |
|  |  |  |  | length | 3583 | 1727 | 1751 |  | 2205 |  |  |  |

kb, kilobases; qtrim, quality trim; hu, human DNA-specific reads; ltrim, length trim.

For error calculation resulting mappings were used. Therefore, the total number of bases as well as the total number of substitutions, insertions and deletions were counted to calculate the respective error rates. Error rate estimation was further divided into the 4 distinct bases in the reference sequence to estimate substitution among bases. Analysis was performed using the function QC for Read Mapping of CLC Genomics Workbench 12.0.

#### Trimming and mapping for ratio estimation

Illumina fastq raw reads were quality trimmed with a base-calling error probability of 0.001, an ambiguous limit of 2 in a read and a minimum and maximum number of nucleotides in reads of 100 and 9000, respectively. Human genomic DNA reads were filtered out as described. First, quality trimmed and filtered reads were randomly mapped to the Merlin and TB40-BAC4-luc amplicon reference sequences, with default setting parameters for match/mismatch scores and insertion/deletion costs, length fraction of 0.3, and similarity fraction of 0.8 to determine the overall % of HCMV-specific reads. Second, for ratio estimation genotype-specific reads mapping to highly polymorphic regions within glycoprotein H (gH) (position in CDS of Merlin strain: 1 to 180), gO (1 to 680), and gN (1 to 220), were considered. In total, 2 gH, 8 gN, and 8 gO genotype sequences were used as reference sequences for mapping (Supplementary Table 6). Mapping parameters were: default setting parameters for match/mismatch scores and insertion/deletion costs, length fraction of 0.3, similarity fraction of 0.95, and non-specific matches were ignored. Merlin and TB40-genotype-specific reads were counted to estimate the ratio. Reads mapping to one or more of the other genotype reference sequences were counted to determine the false-positive mapping rate.

#### Trimming and mapping for genotype determination

Illumina fastq reads were quality trimmed with a base-calling error probability of 0.001, a minimum number of nucleotides in reads of 100, and human genomic DNA reads were removed. Then, quality trimmed and filtered reads were mapped to highly polymorphic regions within gO (1 to 680), gN (1 to 220), and UL146 (position in CDS of Merlin strain: 1 to 360) with default setting parameters for match/mismatch scores and insertion/deletion costs, a length fraction of 0.3 and a similarity fraction of 0.8. A total of 30 genotypic reference sequences were used as previously described [19] and listed in Supplementary Table 6.

A positive genotype mapping was scored if the number of reads was >10 and the consensus length corresponded to at least 80% of the reference length. Consensus sequences were derived from the mappings by using a minimum read depth of 5 reads per base with low coverage regions coded as ambiguities (Ns). Ambiguity codes were inserted using a noise threshold of 0.1 and a minimum nucleotide count of 5. Alignments of derived consensus sequences were screened visually and unique consensus sequences were counted as independent genotypes.

#### Trimming and mapping for haplotype determination

To retain PacBio ccs reads that span the complete amplicon length, all ccs were initially length trimmed. For this, F3 amplicon-derived reads below 7500 and above 8000 and F4 amplicon-derived reads below 6500 and above 7000 number of nucleotides in reads were excluded. Remaining reads were processed with following parameters: quality trimming with a base-calling error probability of 0.01, ambiguous limit of 2 in a read and exclusion of low quality reads. Only reads that span more than half of the amplicon length (4000 nucleotides for F3- and 3500 nucleotides for F4-derived amplicons) were retained and human genomic DNA reads removed. Remaining reads were mapped to 236 HCMV full genomes (Supplementary Table 7), with default setting parameters for match/mismatch scores and insertion/deletion costs, length fraction of 0.7, and similarity fraction of 0.8 and non-specific mappings were ignored. Consensus sequences were generated from mappings with a consensus length of >6.9 kb for F3-and >5.9 kb for F4-derived amplicons, and with a minimum coverage of >0.1% of the average coverage (Supplementary Tables 12a and 12b). Extracted consensus sequences were aligned by muscle [35] and visually inspected in BioEdit 7.2.5 (Ibis Therapeutics). Unique consensus sequences were used as new reference sequences to repeat the mapping with length and similarity fraction of 0.9. New consensus sequences from mappings spanning over the complete amplicon length showing a uniform coverage were extracted. Ambiguity codes were inserted using a noise threshold of 0.1 and a minimum nucleotide count of 5. Unmapped reads were further mapped to human and HCMV genomes to confirm that no haplotype sequences got lost during the analysis steps. After alignment and visual inspection unique consensus sequences were counted as independent haplotypes. To visualise the diversity among distinct haplotypes, nucleotide alignments and phylogenetic tree analysis of all haplotypes were performed in Geneious Prime® 2019.0.3 and Mega 7.0.14 [36], respectively. Phylogenetic trees were inferred by using the Maximum Likelihood method based on the HKY + G (0.05) + I (0,27) for F3 haplotypes and Kimura-2 + G (0,47) + I (0,62) for F4 haplotypes. Best fit substitution model (lowest BIC score) was assessed with Mega 7.0.14.

## Results

### Establishment of long range PCR for highly polymorphic UL regions

First, a long range PCR approach was established by targeting two genomic regions, spanning from UL55 to UL76 (30 kb), and from UL139 to UL146 (6.7 kb), respectively (Figure 1). These two UL regions were primarily chosen based on high interstrain polymorphism and well-characterised genotype assignments of the respective polymorphic genes [19]. The 30 kb long UL region, UL55-UL76, was further divided into 7 segments, which resulted in amplicon sizes ranging from 7.7 kb to 18.0 kb. The target regions exhibit GC contents ranging from 48.5% to 58.9%. Respective primers were designed to align to highly conserved regions (Supplementary Table 1).

To find the best conditions for sensitive and specific long range amplification, three different commercially available PCR enzymes were compared (Supplementary Table 2). Amplification efficiency was tested using highly purified, low-fragmented TB40-BAC4-luc HCMV-DNA isolated from *E.coli*. As shown in Table 1, the enzyme LA Taq Hot Start Version Polymerase delivered the best results in terms of efficiency, fragment length amplification and applicability (2-step PCR).

For further optimization 10-fold dilution series of HFF-derived HCMV-DNA were used, as this source of HCMV-DNA is comparable in composition and integrity to DNA derived from clinical samples. The highest sensitivity was obtained for the shortest amplicons, F3 and F4 (Table 1).

### Error estimation of long amplicon sequencing

Next, long amplicons (>6.6 kb) were sequenced by PacBio and error rates introduced by PCR and sequencing were estimated and compared to Illumina sequencing. To this end, HCMV-DNA of strains TB40-BAC4-luc and Merlin isolated from HFF-infected supernatant were used either as PCR template or directly without any enrichment before sequencing. After generation of ccs and demultiplexing, the resulting reads per sample showed an average length of 3583 to 6162 bases (Table 2). Quality trimming, which was performed with a base-calling error probability (pError) of 0.01 (equals Q>20) and 0.001 (equals Q>30), respectively, resulted in a substantial reduction in read length. This reduction was even more pronounced when raw reads were trimmed to Q>30. Remarkably, up to 19% of all reads had a length <500 bases when quality trimmed to Q>20, and up to 37% when quality trimmed to Q>30. Thus, for subsequent mapping all reads ≤500 bases were excluded after removal of human-specific DNA reads. HCMV-specific mapping rates were ≥85% for non-enriched samples and ≥95% for amplicons. Illumina-derived reads were similarly analysed, except for length trimming of reads <500 bases (Supplementary Table 5).

Error rate was estimated from mapped reads either quality trimmed with pError of 0.01 or 0.001. Comparison between non-enriched and amplicon-enriched DNA allowed to specifically determine the error rates introduced by long range PCR. Substitutions, deletions, and insertions relative to the reference sequence were counted and the percentage of mismatches out of the total number of matched bases was calculated. As shown in Table 3, PacBio sequencing displays a 5.5-fold to 9.5-fold (pError 0.01 and 0.001, respectively) higher substitution error rate for PCR-enriched compared to non-enriched DNA. Indel errors are similarly high in both sample types. Notably, more stringent quality trimming parameters have almost no effect on substitution errors, but can substantially reduce indels. Moreover, insertions are ∼2.0 to 5.0-fold more frequently found than deletions. Taken together, these data show that substitutions are mainly introduced by PCR whereas deletions and insertions mainly result from sequencing. No difference was seen between the two template DNAs, TB40-BAC4-luc and Merlin, and between the distinct target regions for PCR enrichment (Supplementary Table 8).

**Table 3.** Error rates introduced by Long-range PCR and/or sequencing.

| Sequencing platform | Quality trimming | Input DNA | Substitutions* | Deletions* | Insertions* | Substitutions per nucleotide* |  |  |  |
| --- | --- | --- | --- | --- | --- | --- | --- | --- | --- |
|  |  |  |  |  |  | A | C | G | T |
| PacBio | pError 0.01 | amplicons | 0,083 | 0,060 | 0,289 | 0,115 | 0,054 | 0,061 | 0,111 |
|  |  | non-enriched | 0,015 | 0,051 | 0,204 | 0,015 | 0,013 | 0,017 | 0,014 |
| PacBio | pError 0.001 | amplicons | 0,081 | 0,018 | 0,077 | 0,110 | 0,052 | 0,062 | 0,108 |
|  |  | non-enriched | 0,009 | 0,012 | 0,029 | 0,009 | 0,006 | 0,010 | 0,008 |
| Illumina | pError 0.01 | amplicons | 0,163 | 0,005 | 0,002 | 0,242 | 0,096 | 0,089 | 0,240 |
|  |  | non-enriched | 0,083 | 0,004 | 0,001 | 0,091 | 0,077 | 0,078 | 0,090 |
| Illumina | pError 0.001 | amplicons | 0,120 | 0,004 | 0,001 | 0,182 | 0,071 | 0,067 | 0,171 |
|  |  | non-enriched | 0,027 | 0,004 | 0,001 | 0,028 | 0,025 | 0,027 | 0,028 |

\* Error rate is the percentage of mismatches of the total number of bases mapped to the reference sequence. Error rates are means of two distinct input DNAs (Merlin and TB-BAC4-luc, and two amplicons (F1.1+F3, respectively).

Illumina sequencing shows a very low error rate for insertions (0.001%) and deletions (0.004%) whereas substitutions were even 2.0-fold higher compared to PacBio sequencing (Table 3) which could be due to index PCR and bridge amplification steps applied during Illumina library preparation and sequencing.

Finally, a detailed analysis of substitution errors revealed that A and T substitutions are 2.0-fold more frequently found than G and C substitutions (Table 3 and Supplementary Table 9a). This was seen for both sequencing techniques, which further indicates that these substitutions are mainly introduced by PCR (Supplementary Table 9b).

### Ratio estimation of artificial HCMV DNA mixtures after long range PCR

When long amplicon sequencing is intended to be used to estimate the frequency of occurrence of unique haplotypes in mixtures, the PCR enrichment step should guarantee equal amplification of each individual haplotype. For this purpose, purified Merlin and TB40-BAC4-luc HCMV-DNAs, as representatives of two distinct haplotypes, were artificially mixed at defined ratios and subsequently used as template to generate long amplicons (Table 4a). The total amount of template DNA ranged from 4×10^6^ to 9×10^6^ copies per reaction. Amplicons were subjected to Illumina sequencing. Quality trimmed and filtered reads were mapped against a total of 18 unique genotype reference sequences which comprise highly polymorphic regions within gN, gO, and gH (Supplementary Table 6). As seen in Table 4a, the obtained ratios displayed almost the exact same distributions as the original input. This appears to be independent of the input ratios, the polymorphic region tested and the HCMV genome (Supplementary Table 10).

**Table 4a.** Ratio estimation of artificial mixtures used as template DNA for long range PCR.

| **Artificial Mixture** | **HCMV-DNA** | **Aspired input ratio** | **Measured copies/reaction<sup>1</sup>** | **Measured ratio<sup>1</sup>** | **Copy input for PCR** | **Measured ratio<sup>2</sup>** |
| --- | --- | --- | --- | --- | --- | --- |
| 1 | Merlin | 50 | 3,90E+06 | 57 | 3,90E+04 | 44 |
|  | TB40-BAC4-luc | 50 | 2,90E+06 | 43 | 2,90E+04 | 56 |
| 2 | Merlin | 95 | 8,60E+06 | 94 | 8,60E+04 | 94 |
|  | TB40-BAC4-luc | 5 | 4,70E+05 | 6 | 4,70E+03 | 6 |
| 3 | Merlin | 5 | 4,70E+05 | 11 | 4,70E+03 | 3 |
|  | TB40-BAC4-luc | 95 | 3,90E+06 | 89 | 3,90E+04 | 97 |
| 4 | Merlin | 99 | 1,20E+07 | 99 | 1,20E+05 | 98 |
|  | TB40-BAC4-luc | 1 | 1,30E+05 | 1 | 1,30E+03 | 2 |
| 5 | Merlin | 1 | 1,90E+05 | 5 | 1,90E+03 | 1 |
|  | TB40-BAC4-luc | 99 | 3,90E+06 | 95 | 3,90E+04 | 99 |

1) confirmed by HCMV-specific qPCR against gH; 2) measured ratios are means of reads mapping to genotypic regions within gN, gO, and gH; sequencing was done in duplicates.

Next, the lower limit of detection of 1% HCMV DNA in mixtures was determined and TB40-BAC4-luc to Merlin mixtures at a 1:99 ratio were further 10-fold diluted. The 1% HCMV DNA template was detectable down to 19 copies (Table 4b).

**Table 4b.** Ratio estimation of dilution series of artificial mixtures.

| **Artificial Mixture** | **HCMV-DNA** | **Aspired Input ratio** | **estimated copies/reaction** | **Copy input for PCR** | **Measured ratio<sup>2</sup>** |
| --- | --- | --- | --- | --- | --- |
| 1 | Merlin | 1 | 1,90E+05 <sup>1</sup> | 1,90E+05 | 1,5 |
|  | TB40-BAC4-luc | 99 | 3,90E+06 <sup>1</sup> | 3,90E+04 | 98,5 |
| 1:10 | Merlin | 1 | 1,90E+04 | 1,90E+02 | 1 |
|  | TB40-BAC4-luc | 99 | 3,90E+05 | 3,90E+03 | 99 |
| 1:100 | Merlin | 1 | 1,90E+03 | 1,90E+01 | 1,5 |
|  | TB40-BAC4-luc | 99 | 3,90E+04 | 3,90E+02 | 98,5 |
| 1:1000 | Merlin | 1 | 1,90E+02 | 1,90E+00 | 0 |
|  | TB40-BAC4-luc | 99 | 3,90E+03 | 3,90E+01 | 100 |

1) original mixture was confirmed by HCMV-specific qPCR (see Table 4a); 2) measured ratios are means of reads mapping to genotypic regions within gN, gO, and gH.

### Haplotype assessment and linkage analysis in clinical samples with mixed genotype infections

Finally, we aimed to assess whether long read PacBio sequencing is suitable to determine unique haplotypes in clinical samples with mixed HCMV strain infections. Therefore, six BAL samples with a total viral load above 1×10^4^ copies/ml were initially screened for mixed HCMV infections by short amplicon Illumina sequencing. DNA was directly purified from BAL samples and short amplicons targeting highly polymorphic regions within the F3 (gO, gN) and F4 region (UL146) (Figure 2) were generated, pooled and sequenced on an Illumina MiSeq instrument (Supplementary Table 4). Consensus sequences upon mapping against a total of 30 genotype reference sequences were assessed and ambiguity codes included if at least 10% of the reads displayed a variant nucleotide. Samples, BAL2, BAL4, and BAL6 carry two gN, gO, and UL146 genotypes each, and sample BAL5, three distinct gN, gO, and UL146 genotype sequences (Table 5, Supplementary Table 11). These four samples were considered to be mixed HCMV strain infected and further subjected to PacBio long amplicon sequencing.

**Figure 2.**
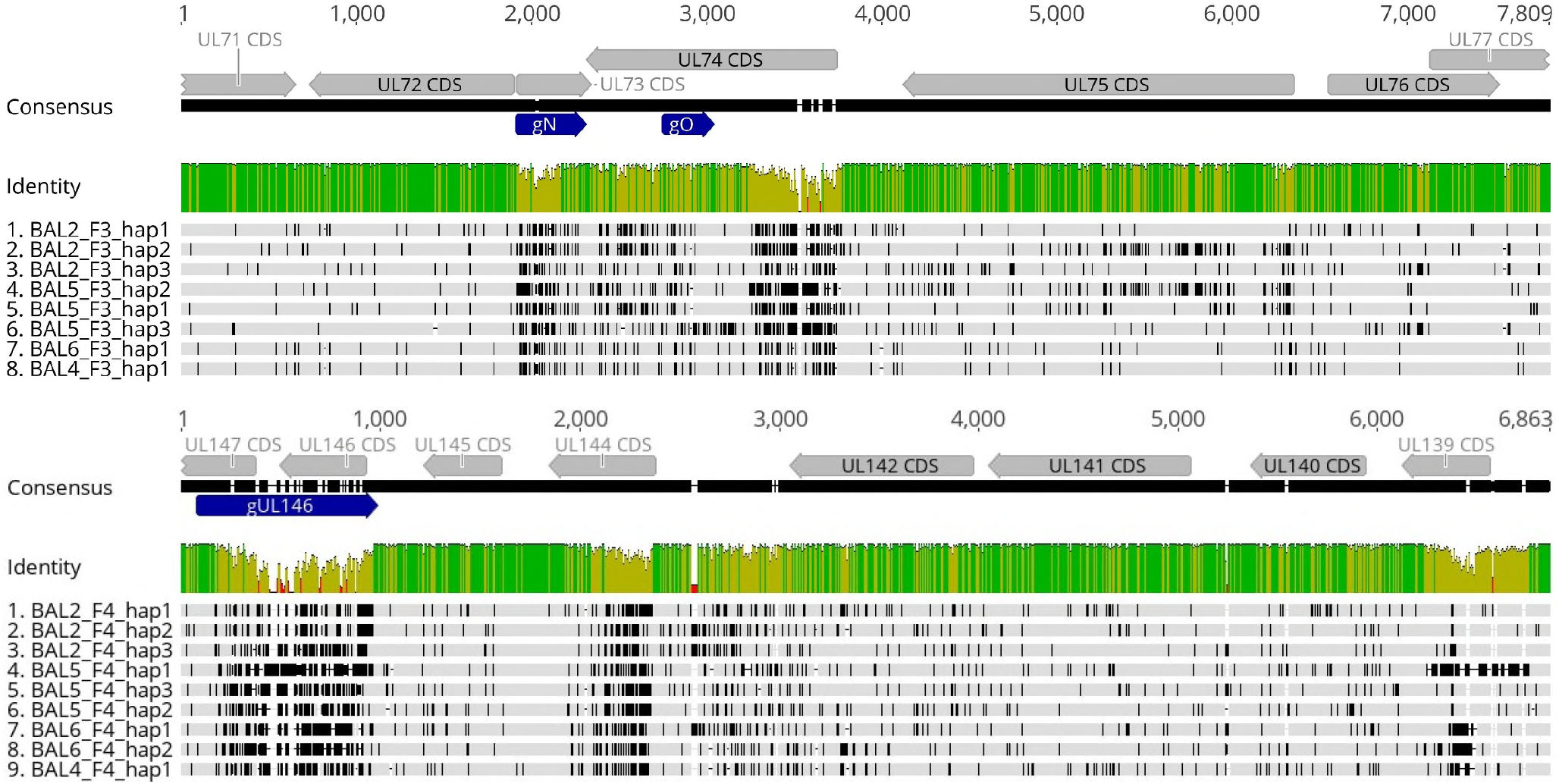
Haplotype diversity of two polymorphic regions, F3 and F4: Shown are sequence alignments of the two polymorphic amplified regions F3 and F4 and their genetic diversity. Top: Haplotype alignments for region F3, Bottom: Haplotype alignments for region F4. Each figure consists of three separate tracks. Alignments of F3 cover a 7.7 kb region (from UL73 to UL76) and F4 alignments cover a 6.7 kb region (from UL146 to UL139). In the top track, annotated CDS and genotypes are depicted with grey or blue arrows, respectively; Track 2: The mean pairwise identity over all pairs in the column is depicted. Green: 100% identity, greenish-brown: at least 30% identity but less than 100%, red: less than 30%; Track 3: Lanes 1-8 or 1-9: Haplotypes sequences, hap, of all bronchoalveolar lavages, BAL, are depicted. Variants are shown in black. Annotations and alignments were generated with geneious prime 2019.0.3.

**Table 5.** Number and ratio of individual haplotype sequences with corresponding gN, gO, and UL146 genotypes as assigned by short amplicon sequencing.

| Sample | Haplotypes determined by long amplicon PacBio sequencing |  |  |  | Genotypes<sup>2</sup> assigned by short amplicon Illumina sequencing |  |  |
| --- | --- | --- | --- | --- | --- | --- | --- |
|  | Target site | Seq ID | Reference sequence<br>Acc.No. (% identity)<sup>1</sup> | Frequency of<br>occurrence (%) | gN | gO | UL146 |
| **BAL2** | F3 | BAL2_F3_hap1 | JX512202.1 (99.9) | 58,7 | 3b | 2a |  |
|  |  | BAL2_F3_hap2 | KP745677.1 (99.1) | 35,1 | 3b | 2a |  |
|  |  | BAL2_F3_hap3 | KJ361966.1 (99.3) | 6,2 | 1 | 1a |  |
|  | F4 | BAL2_F4_hap1 | KP745638.1 (99.9) | 59,2 |  |  | 13 |
|  |  | BAL2_F4_hap2 | KP745654.1 (98.6) | 32,3 |  |  | 13 |
|  |  | BAL2_F4_hap3 | KT726941.2 (98.3) | 8,5 |  |  | 12 |
| **BAL4** | F3 | BAL4_F3_hap1 | KR534203.1 (99.9) | 100 | 1 | 1a |  |
|  |  |  |  |  | 3b | 2a |  |
|  | F4 | BAL4_F4_hap1 | KU550090.1 (99.9) | 100 |  |  | 14 |
|  |  |  |  |  |  |  | 13 |
| **BAL5** | F3 | BAL5_F3_hap1 | KY490065.1 (99.9) | 78,1 | 3b | 2a |  |
|  |  | BAL5_F3_hap2 | JX512208.1 (99.9) | 14,1 | 2 | 2b |  |
|  |  | BAL5_F3_hap3 | KY490066.1 (99.9) | 7,8 | 3a | 1b |  |
|  | F4 | BAL5_F4_hap1 | KR534198.1 (99.8) | 75,1 |  |  | 6 |
|  |  | BAL5_F4_hap2 | KP745691.1 (99.8) | 16,8 |  |  | 7 |
|  |  | BAL5_F4_hap3 | JX512208.1 (99.9) | 8,1 |  |  | 11 |
| **BAL6** | F3 | BAL6_F3_hap1 | KR534203.1 (99.9) | 100 | 1 | 1a |  |
|  |  |  |  |  | 3b | 2a |  |
|  | F4 | BAL6_F4_hap1 | KY490086.1 (99.9) | 85,6 |  |  | 2 |
|  |  | BAL6_F4_hap2 | KY490085.1 (97.1) | 14,4 |  |  | 1 |

1) Sequence in GenBank database with highest similarity to haplotype sequence; 2) according to Suarez et al., 2019, doi: 10.1093/infdis/jiz208; BAL, bronchoalveolar lavage; Acc.No., Accession number.

As listed in Table 5, up to 3 different haplotypes for both, F3 and F4, target sites were identified. These new haplotype sequences have been deposited in Genbank with the accession numbers MW560357-MW560373. Ratio estimation of distinct haplotypes indicates that varying concentrations of haplotypes could be present in mixtures. Notably, BAL2 and BAL5 depict almost the exact same percentage for minor and major haplotypes (Table 5). This could indicate that these non-adjacent haplotype sequences F3 and F4 are located on the same DNA strand of a single strain.

Multiple nucleotide sequence alignments and phylogenetic tree analyses of the 8 unique F3 and the 9 unique F4 haplotype sequences, respectively, were generated (Figure 2 and 3). The overall nucleotide diversity (p-distance) for F3 sequences was 0.045 (range: 0.001 to 0.062) and for F4 was 0.07 (range: 0.034 to 0.099). Sequence comparison of the patient’s haplotype population shows that two sequences can be identical over more than 200 bases and longer stretches of similarity often differ only by a few SNVs (Figure 2). This clearly indicates that short reads are often not long enough to cover non-adjacent diversity. Of note, for all haplotype sequences similar sequences (>97%) were found in the database (Table 5).

**Figure 3.**
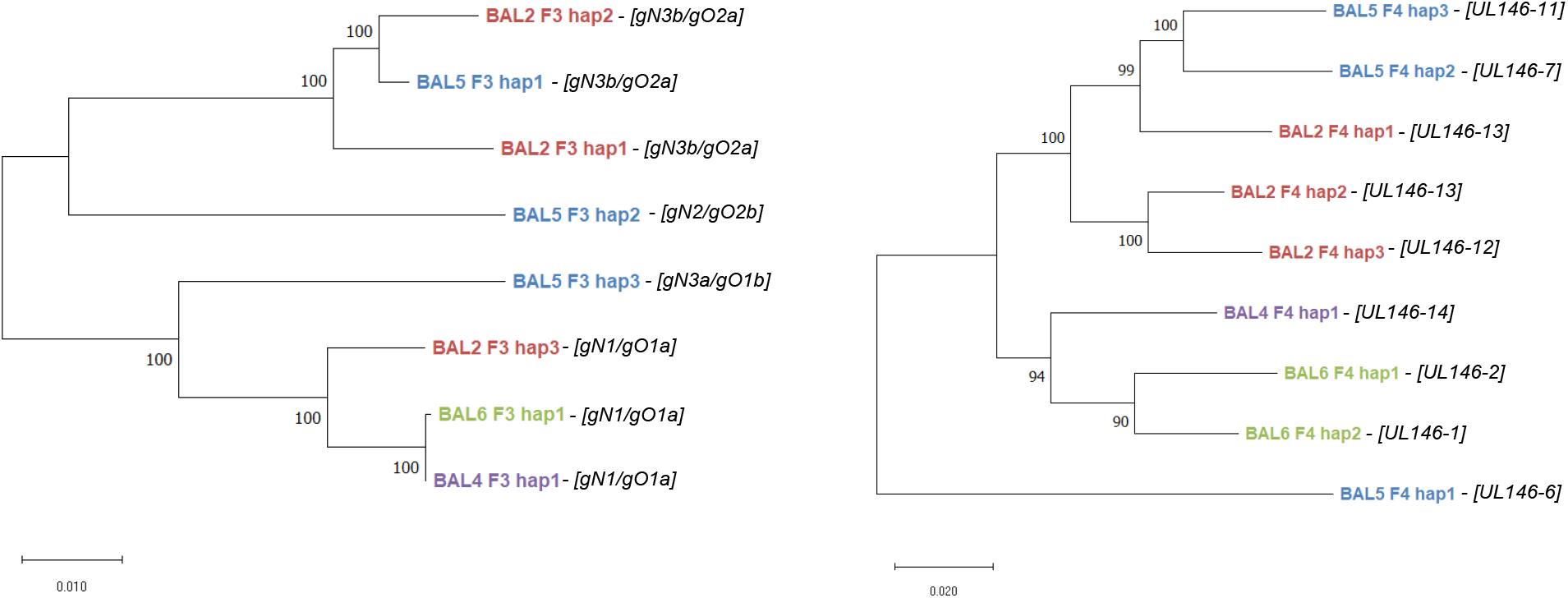
Intra- and interpatient diversity of individual HCMV haplotype sequences. Phylogenetic trees of 8 distinct F3 and 9 distinct F4 haplotype sequences, respectivley, were inferred by using the Maximum Likelihood method and Hasegawa-Kishino-Yano model (F3) or Kimura 2-parameter model (F4). F3 haplotype sequences cover a 7.7kb region (from UL73 to UL76) and F4 haplotype sequences cover a 6.7kb region (from UL146 to UL139) of the HCMV genome. Branches are labeled by the identified haplotypes of the respective sample and amplicon (e.g. BAL2_F3_hap1). Corresponding gN and gO sequences as determined by short amplicon sequencing are shown in bold italics in brackets next to the branch label. BAL, bronchoalveolar lavage; hap, haplotype..

Comparison of both haplotyping and genotyping data revealed that the number of haplotypes and genotypes was numerically the same only for sample BAL5. In BAL4 and BAL6, a higher number of genotypes than haplotypes was detected. These results indicate that low level HCMV DNA sequences are still amplifiable by short range PCR but not by long range PCR. Remarkably, in BAL2 a higher number of haplotypes than genotypes was found. Further inspection of all sequences over the gN, gO, and UL146 region revealed that two distinct F3 haplotype sequences share the same gN3b and gO2a genotype, and two distinct F4 haplotype sequences share the same UL146-13 genotype (Table 5). These shared genotypes, however, vary by a few SNVs. In Figure 4, the data for the two gN3b variants are displayed. Precisely, the two gN3b variants of the respective haplotypes differ by 3 SNVs (p-distance 0.008) over the genotype-defining region whereas the p-distance over the complete F3 amplicon length was 0.026. These findings impressively demonstrate the information benefit of long amplicon over short amplicon sequencing.

**Figure 4:**
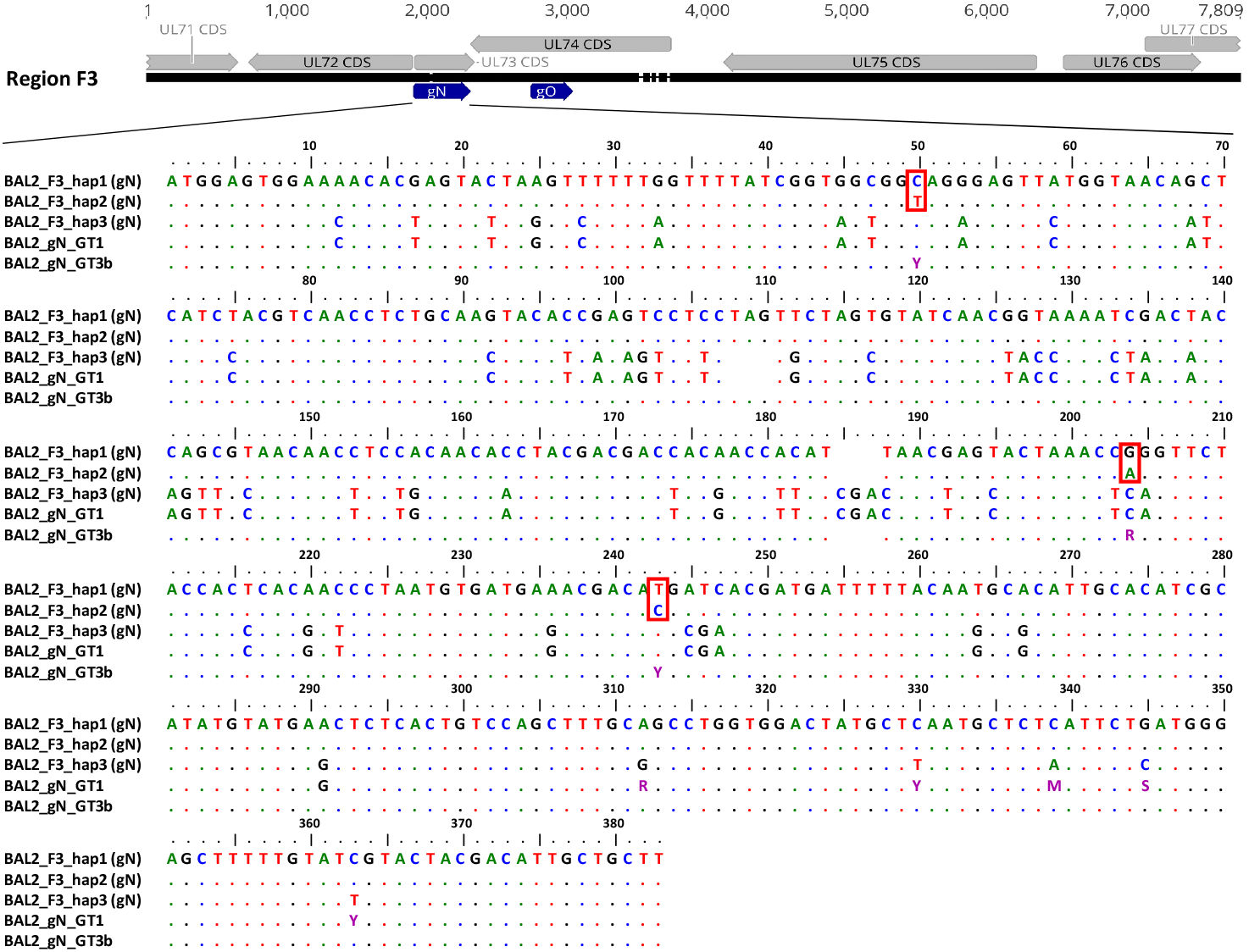
Nucleotide variation between genotype and haplotype sequences of region gN. A depiction of the polymorphic region F3 (7.8 kb) can be seen on the top. While annotated CDS are depicted with grey arrows, genotypes gN (UL73) and gO (UL74), for short read sequencing, can be seen in dark blue. The respective gN region of three haplotype sequences and two gN genotype sequences were aligned. Ambiguous nucleotides are depicted in violet. Red boxes indicate the nucleotide difference between hap 1 and hap 2. Alignments were generated in BioEdit version 7.0.1; GT, genotype; BAL, bronchoalveolar lavage; hap, haplotype.

## Discussion

In this study, we established an amplicon-based deep sequencing approach to determine linkage of interspersed polymorphic regions along the HCMV genome. To the best of our knowledge, this is the first application to characterise and unravel distinct haplotypes within mixed HCMV strain populations by long read sequencing.

### Pros and cons of long range PCR

Initially, we compared different DNA polymerases, all of which were advertised to be suitable for read lengths up to 20 kb. We tested the amplification of 8 HCMV specific target regions, with sizes from 6.7 to 18 kb and observed substantial variations in performance even when highly purified low fragmented HCMV DNA (BAC) was used as template. Irrespective of the average GC content target regions which comprise a GC content of 72% over a 1 kb segment (fragments F2, F2.2, and F1.2) were especially difficult to amplify. Hence, for GC rich regions alternative target enrichment techniques such as solution-based capture [22, 37, 38] and molecular inversion probes [39] might be considered. The enzyme with the best performance was picked out for further optimisation of PCR conditions for the three target regions, F1, F3, and F4, (all <8 kb) using HCMV DNA isolated from cell culture supernatant.

In addition to genome specific limitations, successful amplification of long range amplicons depends on the integrity of input DNA. The extent of DNA fragmentation when directly isolated from patient samples may vary between different sample types and different storage conditions [40]. In this study, we applied our long range PCR protocol on DNA isolated from stored BAL samples. Amplicons with fragment lengths ≤7.7 kb and a GC content ≤54% (F3 and F4) were successfully generated from all samples whereas primer pairs for amplicon F1 (length:.7.8 kb, GC content: 56.5%) failed to amplify PCR products.

Taking these findings together it appears that fragment lengths up to 7 kb and an average GC content of less than 55% are best suited to generate long range amplicons from clinical samples.

### Influence of trimming parameters on read length and error rate

Both, PCR target enrichment and sequencing are prone to false introduction of substitutions and indels [41]. Moreover, it has been reported that PacBio long read sequencing is associated with a higher error rate than short read sequencing techniques [42]. In order to precisely define the error rate of long amplicon sequencing, we compared PCR-enriched with non-enriched samples. Although we observed a substantial reduction in the overall number of reads (25% to 54%) and in the average read length (42% to 72%) depending on the stringency of quality trimming parameters, this was similar for both sample types. Substitution errors, in contrast, were found to be 6-fold to 9-fold higher in amplicons compared to non-enriched samples, yet without any influence of the stringency of quality trimming. These findings suggest that substitution errors are largely introduced by PCR which is well in accordance with previous reports [41]. Overall substitution error rate was about 0.08% which means that low-level single nucleotide variants (SNV) down to 1% could be well distinguishable from PCR-mediated substitutions. SNVs lower than 1%, however, may require additional steps for validation such as performance in duplicates. The number of indels was similar for non-enriched and amplicon-enriched samples indicating introduction by long read sequencing rather than by PCR. For non-ambiguous determination of true insertions and/or deletions either reanalysis with highly quality trimmed reads or an additional short read Illumina sequencing, which shows a very low indel error rate, could be performed. Alternatively, high coverage ccs can be aimed for in order to decrease the error rate down to ∼0.001 % [43]. As can also be seen within our haplotypes, indels are usually the second most abundant form of genetic variation. The reliable detection of indels is still a challenging problem, as our understanding of their origins and functional effects lags behind that of other types of variants. Indels have the potential to generate great changes in a viral population, as truncated or extended genes can be created through frameshifts or by the removal of stop codons. Overall, indels contribute to genetic variation in a virus genome. Within our haplotypes in the subset of samples we did not observe additional stop codons or significant changes in gene length.

When we applied the long read sequencing protocol to determine haplotypes in patient samples we chose a PHRED score of ≥20 for quality trimming. This approach ensures that a higher number of long reads is retained at the expense of a higher indel error rate. Further removal of short reads guarantees that only long reads spanning over multiple non-adjacent polymorphic sites are used for haplotype determination. Depending on the distribution of diversity along the target sites it will be necessary to adapt these analyses parameters accordingly.

### Quantification of haplotype sequences in mixtures

It is critical that each haplotype sequence within a mixture is equally amplified by PCR in order to accurately determine and quantify all haplotypes. For validation, a set of artificial mixtures of two related HCMV DNAs were sequenced. Our data show that the mixed ratios can precisely be determined. Even low abundance haplotypes of 1% were correctly detected down to about 20 copies of the minor haplotype sequence. This finding correlates well with the before mentioned sensitivity limit of about 30 copies per reaction and underlines that both, the total HCMV DNA load and the sensitivity limit of the PCR are critical for detection of minor haplotypes in mixtures, given that the subsequent sequencing step was performed to an appropriate read depth. Since our protocol was validated solely for two haplotypes in mixtures it cannot be excluded that distinct haplotype mixtures or mixtures with more than two haplotypes may influence the PCR performance. All primer pairs, however, were designed to bind to highly conserved regions of the HCMV genome, thus it can be assumed that distinct haplotype sequences are similarly well amplified.

Interestingly, BAL samples which have the same number of haplotypes for regions F3 and F4, displayed almost identical frequencies of occurrence of individual haplotypes for both target sites. It is tempting to speculate that these individual haplotypes are found on the same HCMV DNA molecule within the HCMV population. Such observations may be verified in future studies through a long read based genome sequencing approach with overlapping long amplicons. HCMV displays several diversity hot spots spread across the genome with 30/170 genes having a dN (nonsynonymous substitutions per nonsynonymous site) value of >0.025 [23]. Our two target regions F3 and F4 cover already 7 of these hypervariable genes. To understand the linkage of distantly located genes, however, several overlapping amplicons can be generated.

### Clinical Applicability

Mixed strain infections in transplant recipients are common and have been associated with higher viral loads, delayed viral clearance and poorer clinical outcomes [16]. Potential application of this haplotype determination approach based on long read sequencing in a clinical setting is limited to samples with high viral loads (>10^4^ copies/mL) necessary for long range amplification. We have tested BAL samples for this study, however our lab has successfully used plasma samples as input material as well (data not shown). Since the initial PacBio requirement of a minimum of 700 ng DNA input for library preparation has been reduced to 100 ng, the use of clinical samples with restricted material availability has become even more feasible. Additionally, using multiplex PCRs for long range amplicon generation, patient samples can be used more efficiently. This approach was already tested within our lab and promising results multiplexing F3 and F4 have been achieved.

### Haplotypes versus genotypes

In the present study, the newly developed long read PacBio sequencing approach was applied to four BAL samples that carry multiple genotypes as determined by short amplicon sequencing of gN, gO (located in the F3 target region) and UL146 (located in the F4 target site). Actually, also multiple haplotype sequences were identified by long amplicon sequencing. Comparison of sequencing data from both approaches clearly demonstrates limitations as well as main advantages of long amplicon sequencing. First, our findings indicate a potentially higher detection rate of low abundance populations by short amplicon compared to long amplicon sequencing as observed in two samples. Since short range PCRs usually show a higher sensitivity and are less prone to fragmented DNA found in patient samples, low abundance templates are easier to amplify. This makes this genotyping approach particularly suitable for initial screening of mixed infections. Second, short amplicon genotyping may strongly underestimate the intra-host diversity. Our data show that individual haplotypes can carry the same genotype sequences across the respective target region but are otherwise substantially diverse. Moreover, SNVs of the same genotype which can be detected by short amplicon sequencing will not represent the overall haplotype diversity.

Third, intra-host haplotype sequences show stretches of identity interspersed by diverse sections. This observation underlines the usefulness of long sequencing reads to identify co-linearity of multiple polymorphic regions of individual haplotype sequences which could not be assessed by short reads. Third generation long read sequencing platforms, such as SMRT sequencing from PacBio and Nanopore sequencing from Oxford Nanopore Technologies open this possibility due to its extended read length [28, 29].

Taken together, this study provides the basis of a novel HCMV genome characterization strategy which will lead to an improved understanding of intra-host diversity and the dynamics of mixed HCMV strain populations. This could have important implications for diagnostics, treatment, and vaccine development.

## Abbreviations

BAC: bacterial artificial chromosomes
BAL: bronchoalveolar lavage
css: circular consensus sequence
gH: envelope glycoprotein H
gN: envelope glycoprotein N
gO: envelope glycoprotein O
HCMV: human cytomegalovirus
HFF: human foreskin fibroblast
PacBio: Pacific Biosciences
SMRT: single molecule real-time
SNV: single nucleotide variants
SRA: Sequence Read Archive

## Ethical approval

This pilot study was approved by the Ethics Committee of the Medical University of Vienna under EK-number 1321/2017. All transplant recipients gave their written informed consent and all data were pseudonymised before analyses.

## Acknowledgements

The authors thank Tanja Amon, Andreas Rohorzka, and Ömür Parkan for excellent technical assistance and for their valuable advises on data analysis.

